# Functional normalization of 450k methylation array data improves replication in large cancer studies

**DOI:** 10.1101/002956

**Authors:** Jean-Philippe Fortin, Aurélie Labbe, Mathieu Lemire, Brent W. Zanke, Thomas J. Hudson, Elana J. Fertig, Celia M.T. Greenwood, Kasper D. Hansen

## Abstract

We propose an extension to quantile normalization which removes unwanted technical variation using control probes. We adapt our algorithm, functional normalization, to the Illumina 450k methylation array and address the open problem of normalizing methylation data with global epigenetic changes, such as human cancers. Using datasets from The Cancer Genome Atlas and a large case-control study, we show that our algorithm outperforms all existing normalization methods with respect to replication of results between experiments, and yields robust results even in the presence of batch effects. Functional normalization can be applied to any microarray platform, provided suitable control probes are available.

## Introduction

In humans, DNA methylation is an important epigenetic mark occurring at CpG dinucleotides, implicated in gene silencing. In 2011, Illumina released the IlluminaHuman-Methylation450 bead array [Bibikova et al., 2011], also known as the “450k array”. This array has enabled population-level studies of DNA methylation by providing a cheap, high-throughput and comprehensive assay for DNA methylation. Applications of this array to population level data includes epigenome-wide association studies (EWAS) [Rakyan et al., 2011, Liu et al., 2013] and large-scale cancer studies such as the ones available through The Cancer Genome Atlas (TCGA). To date, around 9000 samples are available from NCBI GEO, and around 8000 samples from TCGA have been profiled on either the 450k array, the 27k array or both.

Studies of DNA methylation in cancer pose a challenging problem for array normalization. It is widely accepted that most cancers show massive changes in their methylome, making the marginal distribution of methylation across the genome different between cancer and normal samples. The global hypomethylation commonly observed in human cancers was recently shown to be organized into large, well-defined domains [Hansen et al., 2011, Berman et al., 2012]. It is worth noting that there are other situations where global methylation differences can be expected, such as between cell types and tissues.

Several methods have been proposed for normalization of the 450k array, including Quantile normalization [Touleimat and Tost, 2012, Aryee et al., 2014], SWAN [Maksimovic et al., 2012], BMIQ [Teschendorff et al., 2013], dasen [Pidsley et al., 2013], and noob [Triche et al., 2013]. A recent review examined the performance of many normalization methods in a setting with global methylation differences and concluded “there is to date no between-array normalization method suited to 450K data that can bring enough benefit to counterbalance the strong impairment of data quality they can cause on some data sets” [Dedeurwaerder et al., 2013]. The authors note that absence of normalization outperforms the methods they evaluate, highlighting the importance of benchmarking any method against raw data.

The difficulties in normalizing DNA methylation data across cancer samples have been recognized for a while. In earlier work on the CHARM platform [Irizarry et al., 2008], Aryee et al. [2011] proposed a variant of subset quantile normalization [Wu and Aryee, 2010] as a solution. The CHARM platform is based on a different assay than the 450k platform. For CHARM, input DNA is compared to DNA processed by a methylation dependent restriction enzyme. Aryee et al. [2011] used subset quantile normalization to normalize the input channels from different arrays to each other. The 450k assay does not involve an input channel; it is based on bisulfite conversion.

Any high-throughout assay suffers from unwanted variation [Leek et al., 2010]. This is commonly addressed by an unsupervised normalization step, followed by application of a batch effect removal tool such as SVA [Leek and Storey, 2007, 2008], ComBat [Johnson et al., 2007] or RUV [Gagnon-Bartsch and Speed, 2012]. The term “batch effect” often refers to unwanted variation remaining after initial normalization, and batch effects have unsurprisingly been observed in studies using the 450K array [Harper et al., 2013].

In this work we propose a method we call functional normalization, which uses control probes to act as surrogates for unwanted variation and some batch effects. We apply this method to the analysis of 450k array data, and show that functional normalization outperforms all existing normalization methods for analysis of datasets with global methylation differences, including studies of human cancer. We also show that functional normalization outperforms the batch removal tool SVA [Leek and Storey, 2007, 2008] in this setting. Our evaluation metrics are focusing on assessing the degree of replication between large-scale studies, arguably the most important, biologically relevant end-point for such studies. Our method is available as the “preprocessFunnorm” function in the minfi package [Aryee et al., 2014] through the Bioconductor project [Gentleman et al., 2004].

### Results

#### Control probes may act as surrogates for batch effects

The 450k array contains 848 control probes. These probes can roughly be divided into negative control probes (613), probes intended for between array normalization (186) and the remainder (51) which are designed for quality control, including assessing the bisulfite conversion rate (see Methods and Supplementary Materials). Important for our proposed method, none of these probes are designed to measure biological signal.

**Figure 1.**
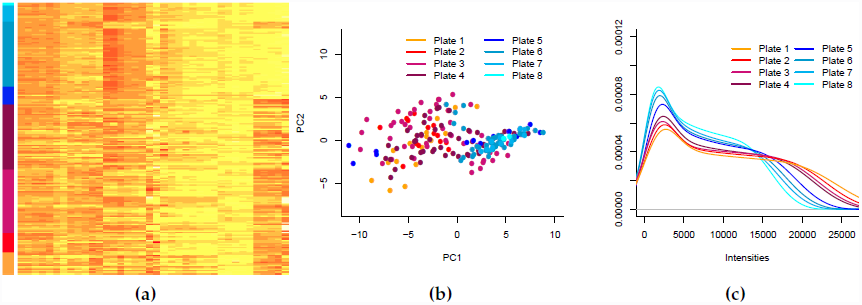
Control probes acts as surrogates for batch effects. (a) A heatmap of a summary (see Methods) of the control probes, with samples on the y-axis and control summaries on the x-axis. Samples were processed on a number of different plates indicated by the color label. Only columns have been clustered. (b) The first two principal components of the matrix depicted in (a). Samples partially cluster according to batch, with some batches showing tight clusters and other being more diffuse. (c) The distribution of methylated intensities averaged by plate. These three panels suggests that the control probe summaries partially measure batch effects.

Figure 1a shows a heatmap of a simple summary (see Methods) of these control probes, for 200 samples assayed on 8 plates (Ontario-EBV dataset). Columns are the control measure summaries and rows are samples. The samples have been processed on different plates, and we observe a clustering pattern correlated with plate. Figure 1b shows the first two principal components of the same summary data and there is evidence of clustering according to plate. Figure 1c shows how the marginal distributions of the methylated channel very across plates. This suggests that the summarized control probes can be used as surrogates for batch effects, as first proposed by Gagnon-Bartsch and Speed [2012] in the context of gene expression microarrays.

#### Functional normalization

We propose functional normalization (see Methods), a method which extends the idea of quantile normalization. Quantile normalization forces the empirical marginal distributions of the samples to be the same, which removes all variation in this statistic. In contrast, functional normalization only removes variation explained by a set of covariates, and is intended to be used when covariates associated with technical variation are available and are independent of biological variation. We adapted functional normalization to data from the 450k array (see Methods), using our observation that the control probe summary measures are associated with technical variability and batch effects. As covariates we recommend using the first *m* = 2 principal components of the control summary matrix, a choice with which we have obtained consistently good results; this is discussed in greater depth below. We have also examined the contributions of the different control summary measures in several different datasets, and we have noted that the control probe summaries given the most weight vary across different data sets.

Functional normalization, like most normalization methods, does not require the analyst to provide information about the experimental design. In contrast, batch removal methods such as SVA [Leek and Storey, 2007, 2008], ComBat [Johnson et al., 2007] and RUV [Gagnon-Bartsch and Speed, 2012] require the user to to provide either batch parameters or an outcome of interest. Like functional normalization, RUV also utilizes the control probes as surrogates for batch effects, but builds the removal of batch effects into a linear model, which returns test statistics for association between probes and phenotype. This limits the use of RUV to a specific statistical model, and methods such as clustering, bumphunting [Aryee et al., 2014, Jaffe et al., 2012] or other regional approaches [Sofer et al., 2013] for identifying differentially methylated regions (DMRs) cannot readily be applied.

#### Funnorm improves the replication between experiments, even when a batch effect is present

As a first demonstration of the performance of our algorithm, we compare lymphocyte samples from the Ontario data set to EBV-transformed lymphocytes samples from the same collection (see Methods). We have recently studied this transformation [Hansen et al., 2014] and have shown that EBV transformation induces large blocks of hypomethylation encompassing more than half the genome, similar to what is observed between most cancers and normal tissue. This introduces a global shift in methylation as shown by the marginal densities in Supplementary Figure S1.

We divided the dataset into discovery and validation cohorts (see Methods), with 50 EBV transformed lymphocytes and 50 normal lymphocytes in each cohort. As illustrated in Supplementary Figure S2a, we introduced an *in silico* batch effect confounding EBV transformation status in the validation cohort (see Methods), to evaluate the performance of normalization methods in the presence of a known confounding batch effect. This has been previously done by others in the context of genomic prediction [Parker and Leek, 2012]. We normalized the discovery cohort, identified the top *k* differentially methylated positions (DMPs) and asked: “how many of these *k* DMPs can be replicated in the validation cohort”. We normalized the validation cohort separately from the discovery cohort to mimic a replication attempt in a separate experiment. We identified DMPs in the validation cohort using the same method and the result is quantified using an ROC curve where the analysis result on the discovery cohort is taken as the gold standard.

**Figure 2.**
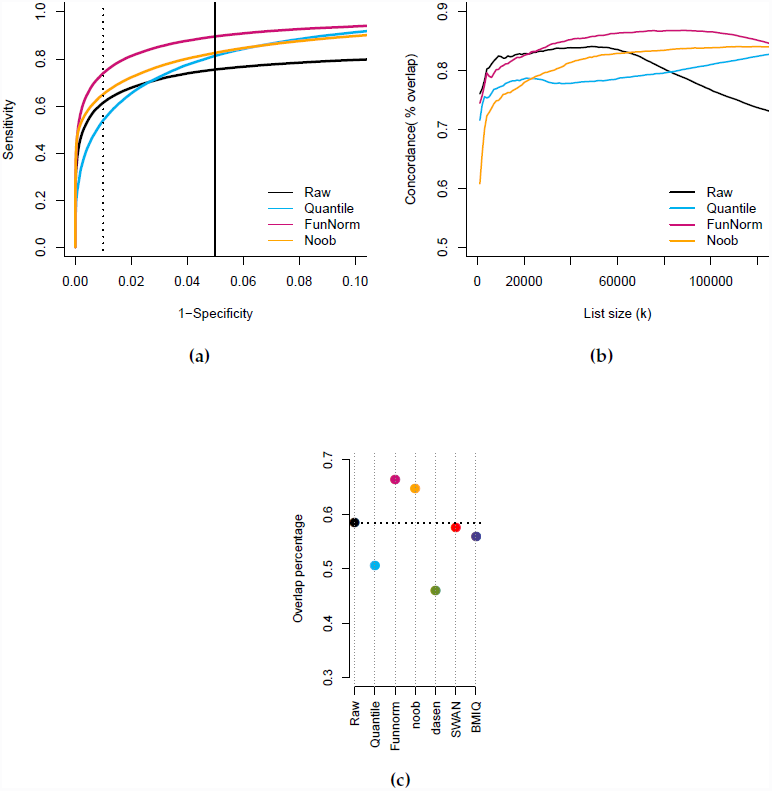
Improvements in replication for the EBV dataset. (a) ROC curves for replication between a discovery and a validation dataset. The validation dataset was constructed to show *in silico* batch effects. The dotted and solid lines represent respectively the commonly used false discovery rate cutoffs of 0.01 and 0.05. (b) Concordance curves showing the percent overlap between the top *k* DMPs in the discovery and validation cohort. Additional normalization methods assessed in Supplementary Figure S3. Functional normalization shows a high degree of concordance between datasets. (c) The percentage of top 100,000 DMPs which are replicated between the discovery and validation cohort and also inside a differentially methylated block or region from Hansen et al. [2014].

To enable the comparison between normalization methods, we fix the number of DMPs across all methods. Because we know from previous work [Hansen et al., 2014] that EBV transformation induces large blocks of hypomethylation covering more than half of the genome, we expected to find a large number of DMPs, and we set *k* = 100, 000. The resulting ROC curves are shown in Figure 2a. In this figure we show, for clarity, what we have found to be the most interesting alternatives to functional normalization in this setting: Raw data, Quantile normalization as suggested by Touleimat and Tost [2012] and implemented in minfi [Aryee et al., 2014] and the noob background correction [Triche et al., 2013]. Supplementary Figures S3a,b contains results for additional preprocessing methods: BMIQ [Teschendorff et al., 2013], SWAN [Maksimovic et al., 2012] and dasen [Pidsley et al., 2013]. Note that each preprocessing method will results in its own set of gold-standard DMPs and these ROC curves therefore measures the internal consistency of each preprocessing method. We note that functional normalization outperforms raw, quantile and noob normalizations when the specificity is above 90% (which is the relevant range for practical use).

We also measured the agreement between the top *k* DMPs from the discovery cohort with the top *k* DMPs from the validation cohort by looking at the overlap percentage. The resulting concordance curves are shown in Figure 2b,with additional methods in Supplementary Figure S3c, and show functional normalization outperforming the other methods.

We can assess the quality of the DMPs replicated between the discovery and validation cohorts by comparing them to the previously identify methylation blocks and differentially methylated regions [Hansen et al., 2014]. In Figure 2c, we present the percentage of the initial *k* = 100, 000 DMPs that are both replicated and present among the latter blocks and regions. We note that these previously reported methylation blocks represent large scale, regional changes in DNA methylation and not regions where every single CpG is differentially methylated. Nevertheless, such regions are enriched for DMPs. This comparison shows that functional normalization achieves a greater overlap with this external dataset, with an overlap of 66% compared to 58% for raw data, while other methods, but noob, perform worse than the raw data.

#### Replication between experiments in cancer study (TCGA-KIRC datasets)

We applied the same discovery-validation scheme to measure performance, used for the analysis of the Ontario-EBV study, on kidney renal clear cell carcinoma samples (KIRC) from TCGA. In total, TCGA has profiled 300 KIRC cancer and 160 normal samples on the 450K platform. Therefore we defined a discovery cohort containing 65 cancers and 65 normals and a validation cohort of 157 cancers and 95 normals (see Methods).

**Figure 3.**
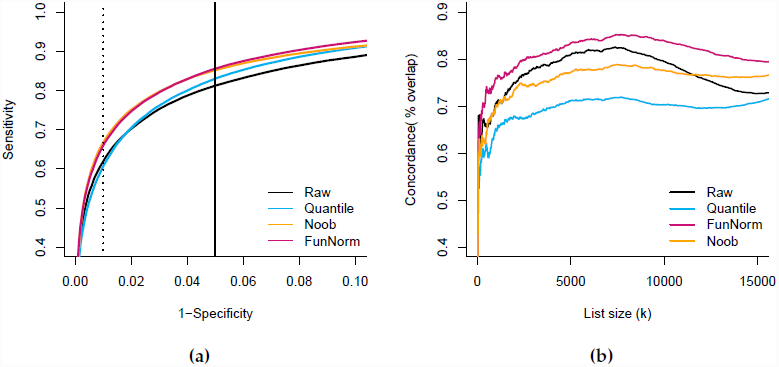
Improvements in replication for the TCGA KIRC dataset. (a) ROC curves for replication between a discovery and a validation dataset. The validation dataset was constructed to show *in silico* batch effects. (b) Concordance plots between an additional cohort assayed on the 27k array and the validation dataset. Additional normalization methods assessed in Supplementary Figure S4. Functional normalization shows a high degree of concordance between datasets.

Our *in silico* batch confounding for this experiment succeeded in producing a validation cohort where the cancer samples have greater variation in background noise (Supplementary Figure S2b). This difference in variation is a less severe batch effect compared to the difference in mean background noise we achieved in the Ontario-EBV dataset (Figure S2a). As for the dataset containing EBV transformed samples, we expect large scale hypomethylation in the cancer samples and therefore we again consider *k* = 100, 000 loci. The resulting ROC curves are shown in Figure 3a, with additional methods in Supplementary Figure S4a,b. Functional normalization and noob are best and do equally well. Again, the gold-standard set of probes that is used to measure performance in these ROC curves differs between normalization methods, and hence these ROC curves reflect the degree of consistency between experiments within each method.

To further compare the quality of the DMPs found by the different methods, we used an additional dataset from TCGA where the same cancer was assayed with the Illumina 27k platform (see Methods). We focused on the 25,978 CpG sites that were assayed on both platforms and asked about the size of the overlap for the top *k* DMPs. For the validation cohort, with maximal batch effects, this is depicted in Figure 3b and Supplementary Figure S4c for additional methods; for the discovery cohort, with minimal batch effects, results are presented in Supplementary Figure S4d. Functional normalization shows the best concordance in the presence of a batch effect in the 450k data (the validation cohort) and is comparable to no normalization in the discovery cohort. Note that while noob and Funnorm show equivalent performance in the ROC curves, only Funnorm improves the validation with an external dataset compared to the raw data.

#### Funnorm preserves subtype heterogeneity in tumor samples (TCGA-AML datasets)

To measure how good our normalization method is at preserving biological variation among heterogeneous samples while removing technical biases, we use 192 acute myeloid leukemia samples from TCGA for which every sample has been assayed on both the 27K and the 450K platforms (see Methods). These two platforms assay 25,978 CpGs in common (but note the probe design changes between array types), and we can therefore assess the degree of agreement between measurements of the same sample on two different platforms, assayed at different time points. The 450k data appears to be affected by batch and dye bias, see Supplementary Figure S5a,b.

**Figure 4.**
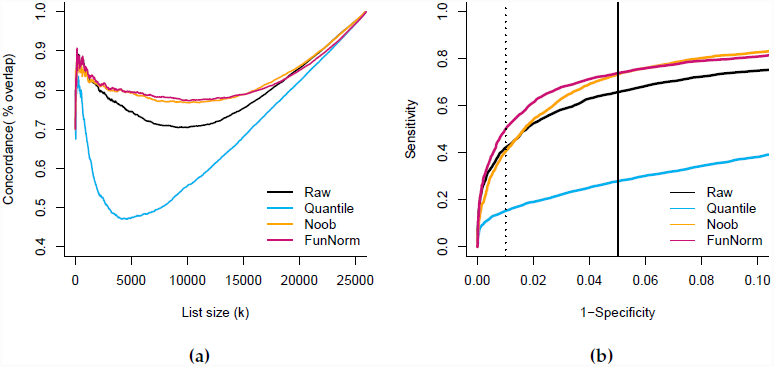
Improvements in replication of tumor subtype heterogeneity. In the AML dataset from TCGA, the same samples have been assayed on 450k and 27k arrays. (a) Concordance plots between results from the 450k array and the 27k array. (b) ROC curves for the 450k data, using the results from the 27k data as gold standard.

Each sample was classified by TCGA according to the French-American-British (FAB) classification scheme [Bennett et al., 1976], which proposes 8 tumor subtypes, and methylation differences can be expected between the subtypes [Figueroa et al., 2010, Akalin et al., 2012]. Using data from the 27k arrays, we identified the top *k* DMPs which distinguish the 8 subtypes. In this case, we are assessing the agreement of subtype variability, as opposed to cancer-normal differences. The analysis of the 27k data uses unnormalized data but adjusts for sample batch in the model (see Methods). Using data from the 450k arrays, we first processed the data using the relevant method, and next identified the top *k* DMPs between the 8 subtypes. The analysis of the 450k data does not include sample batch in the model, which allows us to see how well the different normalization methods remove technical artifacts introduced by batch differences. While both of the analyses are conducted on the full set of CpGs, we focus on the CpGs common between the two platforms and ask “what is the degree of agreement between the top *k* DMPs identified using the two different platforms”. Figure 4a shows that functional normalization and noob outperforms both quantile normalization and raw normalization for all values of *k*, and functional normalization is marginally better than noob for some values of *k*. Supplementary Figure S6a shows the results for additional methods. We can also compare the two datasets using ROC curves, with the results from the 27k data as gold standard (Figure 4b and Supplementary Figure S6b). As DMPs for the 27k data we used the 5,451 CpGs that demonstrate an estimated false-discovery rate less than 5%. On the ROC curve functional normalization and noob outperform Quantile and Raw data for the full range of specificity.

#### Replication between experiments with small changes

**Figure 5.**
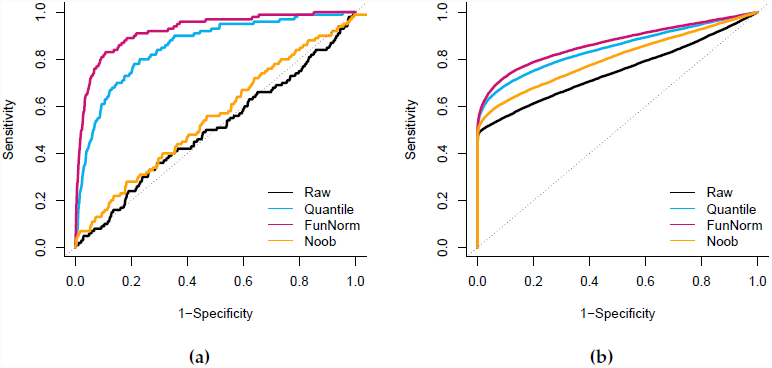
Performance Improvements on blood samples dataset. (a) An ROC curve for replication of case-control differences between blood samples from colon cancer patients and blood samples from normal individuals, the Ontario-Blood dataset. The validation dataset was constructed to show an *in silico* batch effect. (b) An ROC curve for identification of probes on the sex chromosomes for the Ontario-Sex dataset. Sex is confounded by an *in silico* batch effect. Both evaluations show good performance of functional normalization.

To measure the performance of functional normalization in a setting where there are no global changes in methylation, we used the Ontario-Blood dataset which assays lymphocytes from individuals with and without colon cancer. We expect a very small, if any, impact of colon cancer on the blood methylome. As above, we selected cases and controls to form discovery and validation cohorts, and we introduced an *in silico* batch effect that confounds case-control differences in the validation dataset only (see Methods). The discovery and validation datasets contain respectively 283 and 339 samples. For *k* = 100 loci, both functional and quantile normalization show good agreement between discovery and validation datasets, wheres noob and Raw data show an agreement which is not better than a random selection of probes (Figure 5a, Supplementary Figure S8a).

#### Funnorm improves X and Y chromosomes probes prediction in blood samples

As suggested previously [Pidsley et al., 2013], one can benchmark performance by identifying DMPs associated with sex. One copy of the X chromosome is inactivated and methylated in females, and the Y chromosome is absent. On the 450k array, 11,232 and 416 probes are annotated to be on the X and Y chromosomes, respectively. For this analysis it is sensible to remove regions of the autosomes which are similar to the sex chromosomes to avoid artificial false positives that are independent of the normalization step. We therefore remove a set of 30,969 probes which have been shown to cross-hybridize between genomic regions [Chen et al., 2013].

To evaluate our method in the presence of batch effects, we introduced an *in silico* batch effect by selecting 101 males and 105 females from different plates (see Methods), thereby confounding plate with sex. Results show that functional normalization performs well (Figure 5b, Supplementary Figure S8b).

#### Funnorm reduces technical variability

**Figure 6.**
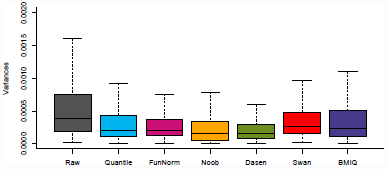
Variance across technical triplicates. Boxplots of the probe-specific variances estimated across 19 individuals assayed in technical triplicates. All normalization methods improve upon Raw data, and Funnorm performs well.

From the Ontario-Replicates lymphocyte dataset (see Methods), we have 19 individuals assayed in technical triplicates dispersed among 51 different chips. To test the performance of each method to remove technical variation, we calculated the probe-specific variance within each triplicate, and averaged the variances across the 19 triplicates. Figure 6 presents box plots of these averaged probe variances of the all methods. All normalization methods improve on Raw data, and functional normalization is in the top 3 of the normalization methods. Dasen in particular does well on this benchmark, which shows that improvements in reducing technical variation do not necessarily lead to similar improvements in the ability to replicate associations.

**Figure 7.**
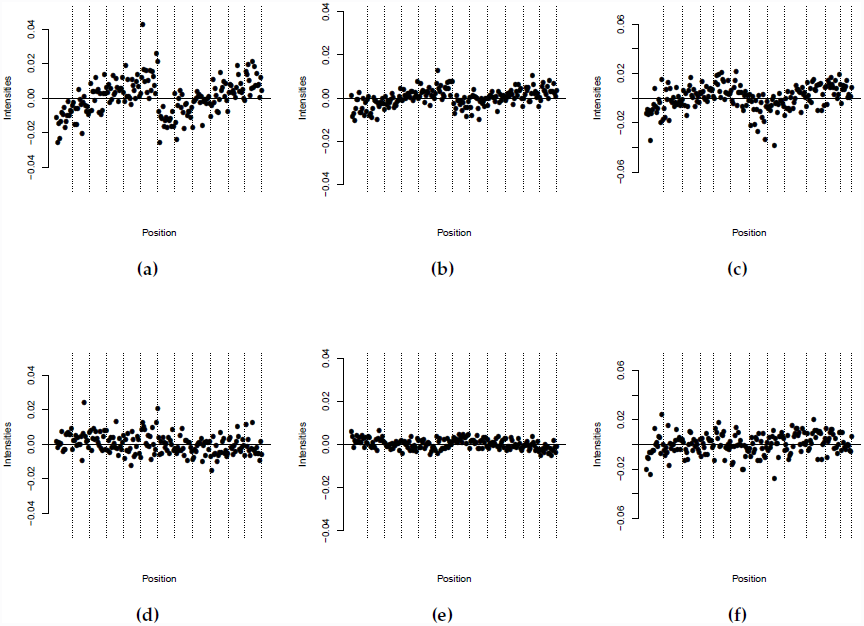
Spatial location affects overall methylation. Quantiles of the beta distributions adjusted for a slide effect. The 12 vertical stripes are ordered as rows 1-6 in column 1 followed by rows 1-6 in column 2. (a) 10*^th^* percentile for Type II probes for the unnormalized AML dataset. (b) 15*^th^* percentile for Type I probes for the unnormalized AML dataset. (c) 85*^th^* percentile for Type II probes for the unnormalized Ontario-Lympho dataset. Panel (a-c) show that the top of the slide has a different beta distribution from the bottom. (d-f) Like (a-c) but after functional normalization, which corrects this spatial artifact.

Each 450k array is part of a slide of 12 arrays, arranged in 2 columns and 6 rows (see Figure 7). Figure 7a–c shows an effect of column and row position on quantiles of the beta value distribution, across several slides. This effect is not present in all quantiles of the beta distribution, and it depends on the dataset which quantiles are affected. Figure 7d–f shows that functional normalization corrects for this spatial artifact.

#### Number of principal components

As described above, we recommend using functional normalization with the number of principal components set to *m* = 2. Supplementary Figure S7 shows the impact of varying the number of principal components on various performance measures we have used throughout, and shows that *m* = 2 always is a good choice. It is outperformed by *m* = 6 in the analysis of the KIRC data and by *m* = 3 in the analysis of the AML data, but these choices perform worse in the analysis of the Ontario-EBV data. While *m* = 2 is a good choice across datasets, we leave *m* to be a user-settable parameter in the implementation of the algorithm. This analysis assumes we use the same *m* for the analysis of both the discovery and validation dataset. We do this to prevent overfitting and to construct an algorithm with no user input. It is possible to obtain better ROC curves by letting the choice of *m* vary between discovery and validation, because one dataset is confounded by batch and the other is not.

#### Comparison to batch effect removal tools

Batch effects are often considered to be unwanted variation remaining after an unsupervised normalization. In the previous assessments we have comprehensively compared functional normalization to existing normalization methods and have shown great performance even in the presence of a batch effect. While functional normalization is an unsupervised normalization procedure, we were interested in comparing its performance to a proper batch removal tool such as SVA [Leek and Storey, 2007, 2008].

**Figure 8.**
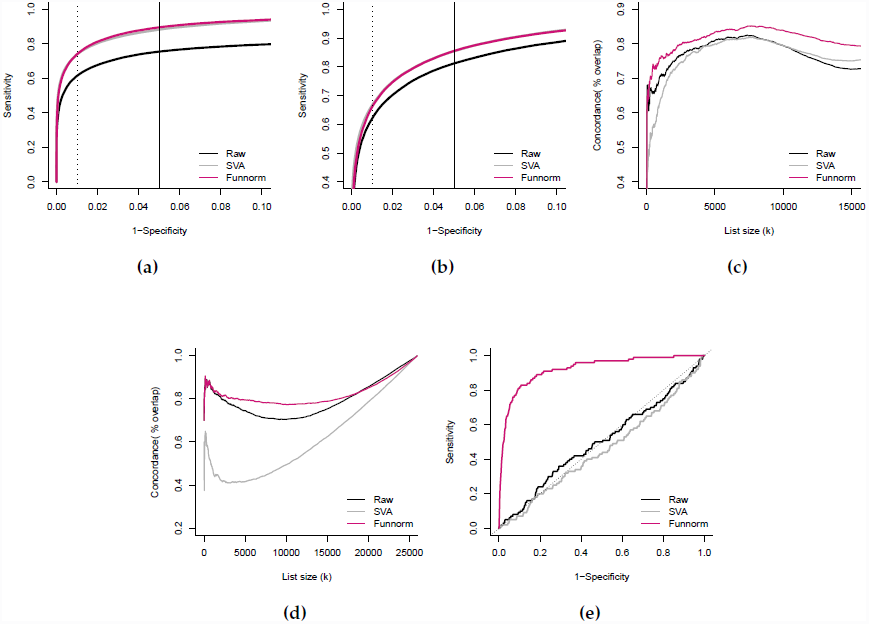
Comparison to batch effect removal tool SVA. (a) Like Figure 2a; an ROC curve for the Ontario-EBV dataset. (b) Like Figure 3a, an ROC curve for the TCGA KIRC dataset. (c) Like Figure 3b, a concordance curve between the validation cohort from 450k data and the 27k data for TCGA KIRC dataset. (d) Like Figure 4a, a concordance plots between results from the 450k array and the 27k array for TCGA AML dataset. (e) Like Figure 5a, an ROC curve for Ontario-Blood dataset.

We applied SVA to all dataset analyzed previously. We let SVA estimate the number of surrogate variables, and allow this estimation to be done separately on the discovery and the validation datasets, which allows for the best possible performance by the algorithm. Figure 8 compares SVA to functional normalization and Raw data for our evaluation datasets. In the analysis of the Ontario-EBV dataset and the TCGA KIRC dataset, SVA and functional normalization achieves similar performance based on the ROC curves, with functional normalization having slightly higher concordance with an external dataset based on the 27k array. For the TCGA AML dataset and the Ontario-Blood dataset functional normalization clearly outperforms SVA.

### Discussion

We have presented functional normalization, an extension of quantile normalization, and have adapted this method to Illumina 450k methylation microarrays. We have shown that this method is especially valuable for normalizing large-scale studies where we expect substantial global differences in methylation, such as in cancer studies or when comparing between tissues, and when the goal is to perform inference at the probe level. Although a normalization method, functional normalization is robust in the presence of a batch effect, and performs better than the batch removal tool SVA on our assessment datasets. This method fills a critical void in the analysis of DNA methylation arrays.

We have evaluated the performance of our method on a number of large scale cancer studies. Critically, we define a successful normalization strategy as one that enhances the ability to reliably detect associations between methylation levels and phenotypes of interest, across multiple experiments. Various other metrics for assessing the performance of normalization methods have been used in the literature on preprocessing methods for Illumina 450k arrays. These metrics include assessing variability between technical replicates [Triche et al., 2013, Teschendorff et al., 2013, Maksimovic et al., 2012, Wu et al., 2013, Dedeurwaerder et al., 2013], and comparing methylation levels to an external gold standard measurement such as bisulfite sequencing [Teschendorff et al., 2013, Dedeurwaerder et al., 2013, Touleimat and Tost, 2012]. We argue that a method which yields unbiased and precise estimates of methylation in a single sample does not necessarily lead to improvements in estimating the differences between samples, yet this is the relevant end-goal for any scientific investigation. For example, it is well known that the RMA method [Irizarry et al., 2003] for analysis of Affymetrix gene expression microarrays introduces bias into the estimation of fold-changes for differentially expressed genes; however, this bias is offset by a massive reduction in variance for non-differentially expressed genes, leading to the method’s proven performance.

In our comparisons, we have separately normalized the discovery and the validation dataset, to mimic replication across different experiments. We have shown that functional normalization is always amongst the top performing methods, whereas other normalization methods tend to perform well on some, but not all, of our test datasets. As suggested by Dedeurwaerder et al. [2013], our benchmarks show the importance of comparing performance to Raw data, which outperforms (using our metrics) some of the existing normalization methods. In several datasets we have observed that the within-array normalization methods, SWAN and BMIQ, have very modest performance compared to Raw data and between-array normalization methods. This suggest that within-array normalization methods does not lead to improvements in the ability to replicate findings between experiments.

Our closest competitor is noob, [Triche et al., 2013], which includes both a background correction and a dye-bias equalization. We outperform noob on our concordance curves in all datasets, and our performance is notably better in the analysis the Ontario-EBV, Ontario-Blood and Ontario-Sex data sets. We find the results on the Ontario-EBV dataset to be particular compelling since the *in silico* batch effect that we introduced was substantial, and yet we were still able to obtain good agreement between discovery and validation analyses.

Our method relies on the fact that control probes carry information about technical batch effects. This idea was previously proposed by Gagnon-Bartsch and Speed [2012] and used to design the batch removal tool RUV. As discussed in the Results section, the RUV method is tightly integrated with a specific statistical model, requires the specification of the experimental design, and cannot readily accommodate regional methods [Aryee et al., 2014, Jaffe et al., 2012, Sofer et al., 2013] nor clustering. In contrast, functional normalization does not require specification of the experimental design and is applied as a preprocessing step. Batch effects are often considered to be unwanted variation remaining after an unsupervised normalization, and we conclude that functional normalization removes a greater amount of unwanted variation in the preprocessing step.

However, control probes cannot measure unwanted variation arising from factors such as cell type heterogeneity, which is known to be an important confounder in methylation studies of samples containing mixtures of cell types [Jaffe and Irizarry, 2014]. This is an example of unwanted variation from a biological, as opposed to technical, source. Cell type heterogeneity is a particular challenge in EWAS studies of whole blood, but this requires other tools and approaches to address.

Surprisingly, we show that functional normalization improves on the batch removal tool SVA applied to raw data, in the datasets we have assessed. It is a very strong result that an unsupervised normalization method improves on batch removal tool, which requires the specification of the comparison of interest.

While we have shown that functional normalization performs well in the presence of batch effects, we still recommend that any large-scale study consider the application of batch removal tools such as SVA [Leek and Storey, 2007, 2008], ComBat [Johnson et al., 2007] or RUV [Gagnon-Bartsch and Speed, 2012] after using functional normalization, due to their proven performance and their potential for removing unwanted variation which cannot be measured by control probes. As an example, Jaffe and Irizarry [2014] discusses the use of tools to control for cell type heterogeneity.

The analysis of the Ontario-Blood dataset suggests that functional normalization has potential to aid the analysis in a standard EWAS setting, where only a small number of differentially methylated loci are expected. However, if only very few probes are expected to demonstrate changes, and if those changes are small, it becomes difficult to evaluate the performance of our normalization method using our criteria of successful replication.

The main ideas of functional normalization can readily be applied to other microarray platforms, including gene expression and miRNA arrays, provided that the platform of interest contains a suitable set of control probes. We expect the method to be particular useful when applied to data with large anticipated differences between samples.

## Acknowledgements

### Funding

Funding for the methylation genotyping was obtained from a GL2 grant from the Ontario Research Fund to BWZ and TJH. CCFR is supported by the National Cancer Institute, National Institutes of Health under RFA #CA-95-011. The OFCCR is supported by grant U01 CA074783. JPF was partially supported by les Fonds de Recherche en Santé du Québec, awarded to AL, the Natural Sciences and Engineering Research Council of Canada and by les Fonds de recherche Nature et technologies du Québec, and a pilot project from the Johns Hopkins Head and Neck Cancer SPORE awarded to EJF. Research reported in this publication was supported by National Institute of General Medical Sciences of the National Institutes of Health under award number R01GM083084. TJH and BWZ are recipients of Senior Investigator Awards from the Ontario Institute for Cancer Research, through generous support from the Ontario Ministry of Research and Innovation. EJF is also supported by the National Cancer Institute (CA141053).

The content of this manuscript does not necessarily reflect the views or policies of the National Cancer Institute or any of the collaborating centers in the CFRs, nor does mention of trade names, commercial products or organizations imply endorsement by the U.S. Government or the CFR. The content is solely the responsibility of the authors and does not necessarily represent the official views of the National Institutes of Health.

*Conflict of Interest:* None declared.

## Methods

### Infinium HumanMethylation450 BeadChip

We use the following terminology, consistent with the minfi package [Aryee et al., 2014]: the 450k array is available as slides consisting of 12 arrays. These arrays are arranged in a 6 row by 2 columns layout. The scanner can process up to 8 slides in a single plate.

We use the standard formula *β* = *M*/(*M* + *U* + 100) for estimating percent methylation given (un)methylation intensities *U* and *M*.

Functional normalization uses information from the 848 control probes on the 450k array, as well as the “out-of-band” probes discussed in Triche et al. [2013]. These control probes are not part of the standard output from GenomeStudio, the default Illumina software. Instead we use the IDAT files from the scanner together with the open source illuminaio [Smith et al., 2013] package to access the full data from the IDAT files. This step is implemented in minfi [Aryee et al., 2014].

### Control probe summaries

We transform the 848 control probes, as well as the out-of-band probes [Triche et al., 2013] into 42 summary measures. The control probes contributes 38 of these 42 measures and the out-of-band contributes 4. An example of a control probe summary is the mapping of 61 “C” normalization probes to a single summary values, their mean. The out-of-band probes are the intensities of the Type I probes measured in the opposite color channel from the probe design. For the 450k platform, this means 92,596 green intensities, and 178,406 red intensities that can be used to estimate background intensity, and we summaries these values into 4 summary measures. A full description of how the control probes and the out-of-band probes are transformed into the summary control measures are listed in the Supplementary Material.

### Functional normalization: the general framework

Functional normalization extends the idea of quantile normalization, by adjusting for known covariates measuring unwanted variation. In this section we present a general model that is not specific to methylation data. The adaptation of this general model to the 450k data is discussed in the next section. The general model is as follows. Consider **Y**_1_,…, **Y**_*n*_ independent high dimensional vectors each associated with a set of scalar covariates *Z_i,j_* with *i* = 1,…, *n* indexing samples and *j* = 1,…,*m* indexing covariates. Ideally these known covariates are associated with unwanted variation and unassociated with biological variation; functional normalization attempts to remove their influence.

For each high-dimensional observation **Y**_*i*_, we form the empirical quantile function for its marginal distribution, and denote it by 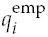 Quantile functions are defined on the unit interval and we use the variable *r* ∈ [0,1] to evaluate them pointwise, like 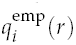. We assume the following model in pointwise form

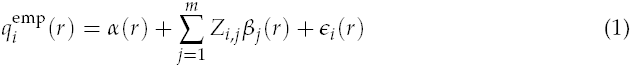

which has the functional form

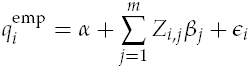

The parameter function *α* is the mean of the quantile functions across all samples, *β_j_* are the coefficient functions associated with the covariates and *∊_i_* are the error functions which are assumed to be independent and centered around 0.

In this model, the term

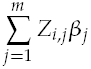

represents variation in the quantile functions explained by the covariates. By specifying known covariates that measure unwanted variation and that are not associated with biological signal, functional normalization removes unwanted variation by regressing out the latter term. An example of a known covariate could be processing batch. In a good experimental design, processing batch will not be associated with biological signal.

In particular, assuming we have obtained estimates 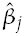 for *j* = 1,…, *m*, we form the functional normalized quantiles by

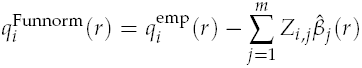

We then transform **Y**_*i*_ into the functional normalized quantity 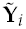 using the formula

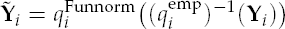

This ensures that the marginal distribution of 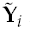 has 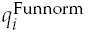 as its quantile function.

We now describe how to obtain estimates 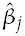 for *j* = 1,…, *m*. Our model 1 is an example of function-on-scalar regression, described in [Reiss et al., 2010]. The literature on function-on-scalar regression makes assumptions about the smoothness of the coefficient functions and uses a penalized framework because the observations appear noisy and non-smooth. In contrast, because our observations **Y**_*i*_ are high-dimensional, the observed empirical quantile functions have a high degree of smoothness. This allows us to circumvent the smoothing approach used in traditional function-on-scalar regression. We use a dense grid of *H* equidistant points between 0 and 1, and we assume that *H* is much smaller than the dimension of **Y**_*i*_. On this grid, model 1 reduces pointwise to a standard linear model. Because the empirical quantile functions *q*^emp^ (*r*) are smooth in *r*, the parameter estimates are smooth across *r* as well. This allows us to use *H* standard linear model fits to compute estimates 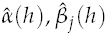 *j* = 1,…, *m* with *h* being on the dense grid {*h* ∈ *d/H*: *d* = 0,1,…, *H*}. We next form estimates 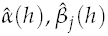 *j* = 1,…, *m* for any *r* ∈ [0,1] by linear interpolation. This is much faster than traditional approaches in the function-on-scalar regression literature.

Importantly, in this framework, using a saturated model in which all the variation (but the mean) is explained by the covariates results in removing all variation and is equivalent to quantile normalization. In our notation, quantile normalized quantile functions are

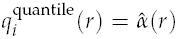

where 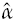 is the mean of the empirical quantile functions. This corresponds to the maximum variation that can be removed in our model. In contrast, including no covariates makes the model comparable to no normalization at all. By choosing covariates which only measure unwanted technical variation, functional normalization will only remove the variation explained by these technical measurements and will leave biological variation intact. Functional normalization allows a sensible tradeoff between not removing any technical variation at all (no normalization) and removing too much variation, including global biological variation, as it can occur in quantile normalization.

### Functional normalization for 450k arrays

We apply the functional normalization model to the methylated (M) and unmethylated (U) channels separately. Since we expect the relationship between the methylation values and the control probes to differ between Type I and Type II probes, functional normalization is also applied separately by probe type to obtain more representative quantile distributions. We address probes mapping to the sex chromosomes separately, see below. This results in 4 separate applications of functional normalization, using the exact same covariate matrix, with more than 100,000 probes in each normalization fit. For functional normalization, we pick *H* = 500 equidistant points (see notation in previous section). As covariates we use the first *m* = 2 principal components of the summary control measures as described above. We do this because the control probes are not intended to measure biological signal since they are not designed to hybridize to genomic DNA. Our choice of *m* = 2 is based on empirical observations on several datasets.

Following the ideas from quantile normalization for 450k arrays Touleimat and Tost [2012], Aryee et al. [2014], we normalize the probes mapping to the sex chromosomes (11,232 and 416 probes for the X and Y chromosomes respectively) separately from the autosomal probes. For each of the two sex chromosomes, we normalize males and females separately. For the X chromosome we use functional normalization and for the Y chromosome we use quantile normalization since the small number of probes on this chromosome violates the assumptions of functional normalization which results in instability.

Functional normalization only removes variation in the marginal distributions of the two methylation channels associated with control probes. This preserves any biological global methylation difference between samples.

### Data

#### The Ontario study

The Ontario study consists of samples from 2200 individuals from the Ontario Familial Colon Cancer Registry (OFCCR) [Cotterchio et al., 2000] who had previously been genotyped in a case-control study of colorectal cancer in Ontario [Zanke et al., 2007]. The majority of these samples are lymphocytes derived from whole blood. We use various subsets of this dataset for different purposes.

**The Ontario-EBV dataset.** Lymphocyte samples from 100 individuals from the Ontario study were transformed into immortal lymphoblastoid cell lines (LCL) using EBV transformation. We divided the 100 EBV-transformed samples into two equal equally sized datasets (discovery and validation). For the discovery dataset, we matched the 50 EBV-transformed samples to 50 other lymphocyte samples assayed on the same plates. For the validation dataset, we matched the 50 EBV-transformed samples to 50 other lymphocyte samples assayed on different plates.

**The Ontario-Blood dataset.** From the Ontario study, we first created a discovery-validation design where we expect only a small number of loci to be differentially methylated. For the discovery dataset, we selected all cases and controls on 3 plates showing little evidence of plate effects among the control probes, which yielded a total of 52 cases and 231 controls. For the validation dataset, we selected 4 plates where the control probes did show evidence of a plate effect and then selected cases and controls from separate plates, to maximize a confounding effect of plate. This yielded a total of 175 cases and 163 controls.

**The Ontario-Sex dataset.** Among 10 plates for which the control probes demonstrated differences in distribution depending on plate, we selected 101 males from a set of 5 plates and 105 females from another set of 5 plates, attempting to maximize the confounding effect of batch on sex.

**The Ontario-Replicates dataset.** Amongst the lymphocyte samples from the Ontario study, 19 samples have been assayed 3 times each. One replicate is a hybridization replicate and the other replicate is a bisulfite conversion replicate. The 57 samples have been assayed on 51 different slides across 11 different plates.

**The TCGA-KIRC datasets.** From The Cancer Genome Atlas (TCGA) we have access to kidney renal clear cell carcinoma and normal samples, assayed on two different methylation platforms. We use the level 1 data which contains the IDAT files. For the 450k platform, TCGA has assayed 300 tumor samples and 160 normal samples. For the discovery set, we select 65 tumor samples and 65 matched normals from slides showing little variation in the control probes. These 130 samples were assayed on 3 plates. For the validation dataset we select the remaining 95 normal samples together with all 157 cancer samples part of the same TCGA batches as the 95 normals. These samples were spread over all 9 plates, therefore maximizing potential batch effects. For the 27k platform, TCGA has assayed 219 tumor samples and 199 normals. There is no overlap between the individuals assayed on the 450k platform and the individuals assayed on the 27k platform.

**The TCGA-AML datasets.** Also from TCGA, we used data from 194 acute myeloid leukemia (AML) samples, where each sample was assayed twice: first on the 27K Illumina array and subsequently on the 450K array. Every sample but 2 have been classified according to the French-American-British (FAB) subtype classification scheme [Bennett et al., 1976] which classifies the tumor into one of 8 subtypes. The 2 unclassified samples were removed post-normalization. We use the data known as level 1, which contains the IDAT files.

**WGBS EBV data**. Hypomethylated blocks and small differentially methylated regions (DMRs) between transformed and quiescent cells were obtained from a previous study [Hansen et al., 2014]. Only blocks and DMRs with a family-wise error rate equal to 0 was retained (see the reference). A total of 228,696 probes on the 450K array overlap with the blocks and DMRs.

### Comparison to normalization methods

We have compared our functional normalization, “Funnorm”, to the most popular normalization methods used for the 450k array. This includes the following between-array normalization methods: (1) “Quantile”: stratified quantile normalization as proposed by Touleimat and Tost [2012] and implemented in minfi [Aryee et al., 2014], (2) “dasen”: background adjustment and between-sample quantile normalization of M and U separately as implemented in Pidsley et al. [2013] and (3) “noob”: a background adjustment model using the out-of-band control probes followed by a dye bias correction [Triche et al., 2013], implemented in the methylumi package. We also consider two within -array normalization methods: (4) “SWAN”: Subset-quantile within-array normalization [Maksimovic et al., 2012] and (5) “BMIQ”: Beta-mixture quantile normalization [Teschendorff et al., 2013]. Finally, we consider (6) “Raw” data: no normalization.

In its current implementation, noob yields missing values for at most a couple of thousand loci per array. This is based on excluding loci below an array-specific detection limit. We have discarded those loci from our performance measures, but only for the noob performance measures. In its current implementation, BMIQ produced missing values for all type II probes in 5 samples for the TCGA AML dataset. We have excluded these samples for our performance measures, but only for our BMIQ performance measures.

For clarity, in figures we focus on the top-performing methods which are Quantile, Raw and noob. The assessments of the other methods, dasen, BMIQ and SWAN are available in Supplementary Materials.

### Comparison to batch removal tools

We used the reference implementation of SVA in the “sva” package [Leek et al., 2012]. We applied SVA to the M-values obtained from the raw data. Surrogate variables were estimated using the iteratively re-weighted surrogate variable analysis algorithm [Leek and Storey, 2008], and were estimated separately for the discovery and validation cohorts. In the analysis of the Ontario-EBV dataset, SVA found 21 and 24 surrogate variables respectively for the discovery and the validation cohorts. In the analysis of the Ontario-Blood dataset, SVA found 18 and 21 surrogate variables respectively for the discovery and the validation cohorts. In the analysis of the TCGA KIRC dataset, SVA found 29 and 32 surrogate variables respectively for the discovery and the validation cohorts. In the analysis of the TCGA LAML dataset, SVA found 24 surrogate variables.

### Identification of differentially methylated positions

To identify differentially methylated positions (DMPs), we used F-statistics from a linear model on the beta values from the array. The linear model was applied on a probe-byprobe basis. In most cases, the model includes case/control status as a factor. In the 27K data, we adjust for batch by including a plate indicator (given by TCGA), as a factor in the model.

### Discovery-Validation comparisons

To measure the consistency of each normalization method at finding true DMPs, we compare results obtained on a discovery-validation split of a large dataset. Comparing results between two different subsets of a large dataset is an established idea and have been applied to the context of 450k normalization [Wu et al., 2013, Teschendorff et al., 2013]. We extend this basic idea in a novel way by introducing an *in silico* confounding of treatment (case/control status) by batch effects as follows. In a first step, we selected a set of samples to be the discovery cohort, by choosing samples where the treatment variable is not visibly confounded by plate effects. Then the validation step is achieved by selecting samples demonstrating strong potential for treatment confounding by batch, for example by choose sample from different plates (see Data descriptions). The extent to which it is possible to introduce such a confounding is dataset dependent. In contrast to earlier work Wu et al. [2013], we normalize the discovery and the validation cohort separate, to more realistic mimic an independent replication experiment. The idea of creating *in silico* confounding between batch and treatment has been previously explored in the context of genomic prediction [Parker and Leek, 2012].

We quantify the agreement between validation and discovery in two ways: by an ROC curve and a concordance curve.

For the ROC curve we use the discovery cohort as the gold-standard. Because the validation cohort is affected by a batch effect, a normalization method that is robust to batch effects will show better performance in the ROC curve. Making this ROC curve requires us to choose a set of DMPs for the discovery cohort. The advantage of the ROC curve is that the plot displays immediately interpretable quantities such as specificity and sensitivity.

For the concordance curve we compare the top *k* DMPs from the discovery and the validation set each, and display the percentage of the overlap for each *k*. These curves do not require us to select a set of DMPs for the discovery cohort. Note that these curves have been previously used in the context of multiple-laboratory comparison of microarray data [Irizarry et al., 2005].

## Supplementary Figures

**Supplementary Figure S1.**
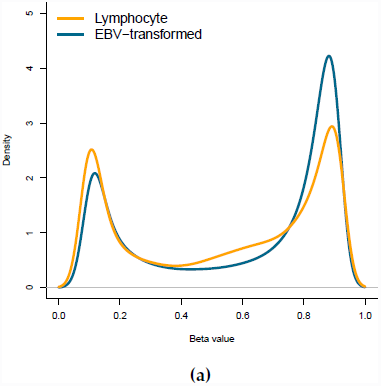
Global changes in DNA methylation for the EBV dataset. (a) The average density of unnormalized beta values across both EBV transformed lymphocytes and normal lymphocytes, showing global hypomethylation caused by EBV transformation.

**Supplementary Figure S2.**
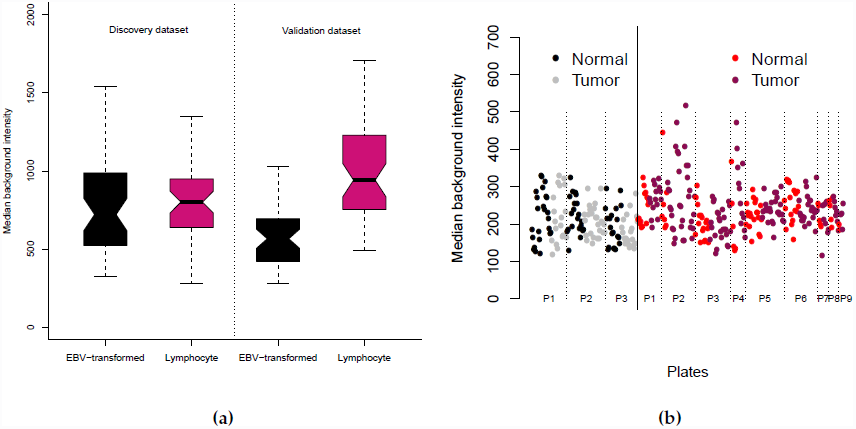
Illustration of *in silico* batch effects. (a) Distribution of background intensity for the Ontario-EBV dataset, showing an *in silico* introduced difference in the validation dataset. (b) Distribution of background intensity for the TCGA KIRC dataset, showing an *in silico* introduced difference in variation in the validation dataset.

**Supplementary Figure S3.**
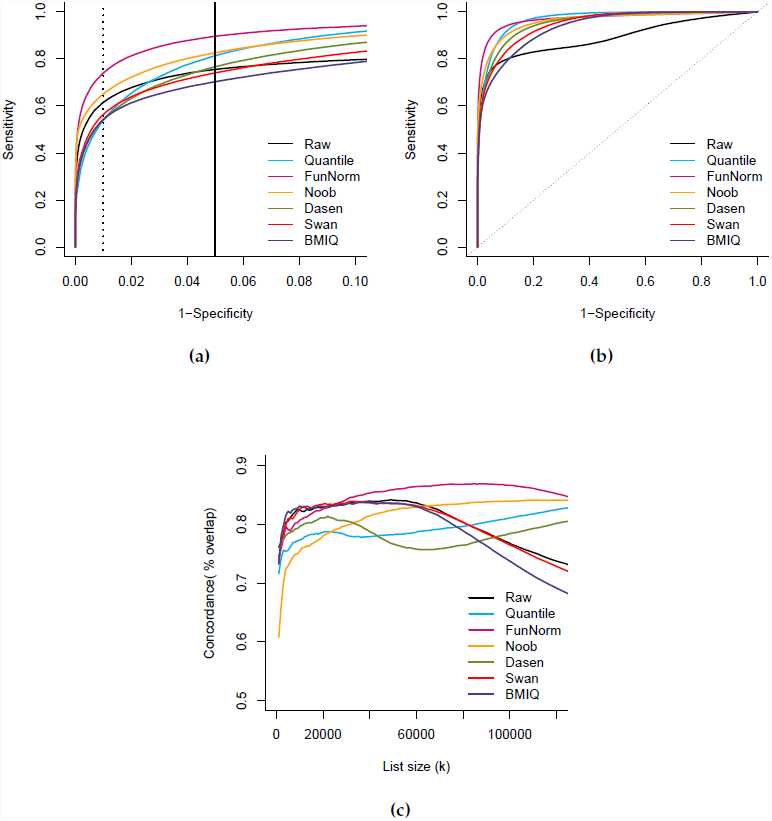
Improvements in replication for the EBV dataset, all methods. (a) Like Figure 2b, but for all examined normalization methods. (b) Like (a) but for the full range of specificity. (c) Like Figure 2c but for all examined normalization methods.

**Supplementary Figure S4.**
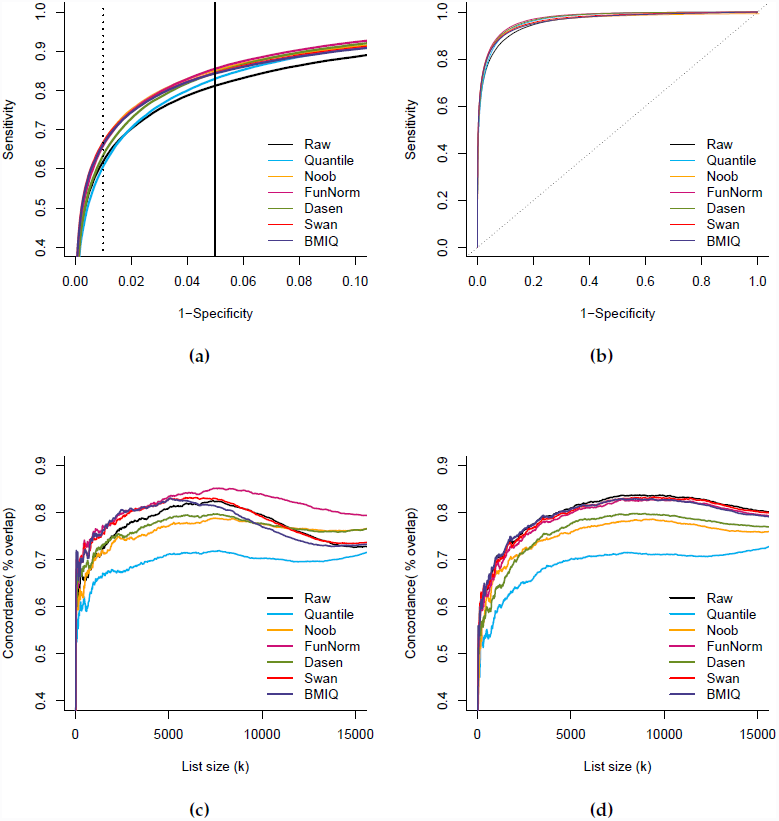
Improvements in replication for the TCGA KIRC dataset, all methods. (a) Like Figure 3a but for all normalization methods we assess. (b) Like (a) but for the full range of specificity. (c) Like Figure 3b but for all normalization methods we assess. (d) Like (c) but comparing to the discovery dataset instead of the validation dataset; the discovery dataset is less affected by unwanted variation.

**Supplementary Figure S5.**
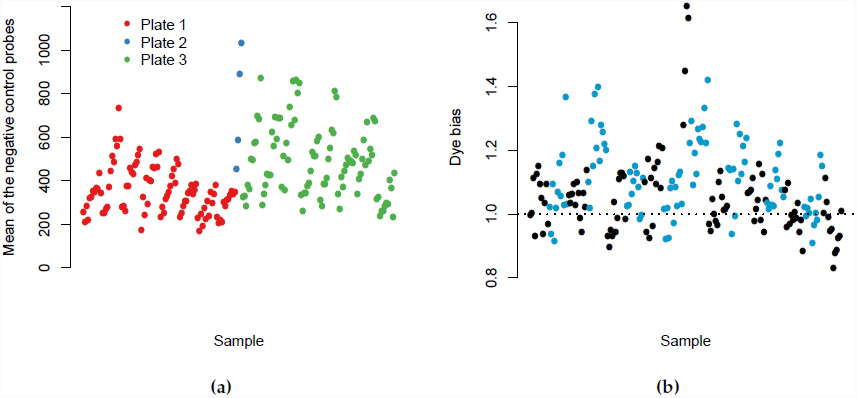
Plate effects and dye bias for the AML dataset. (a) The means of the negative control probes is correlated with the processing plate (96 samples) indicating that background intensity is affected by batch. (b) We measure dye bias by taking the ratio of the negative control probes in the green channel and the negative control probes in the red channel. A value of 1 means that there is no dye-bias. We plot the dye bias (y-axis) for samples ordered by plate and then by slide (x-axis). The plate order is the same as (a). We use two different alternating colors to differentiate the slides. We observe that dye bias is orthogonal to plate effect and highly slide dependent.

**Supplementary Figure S6.**
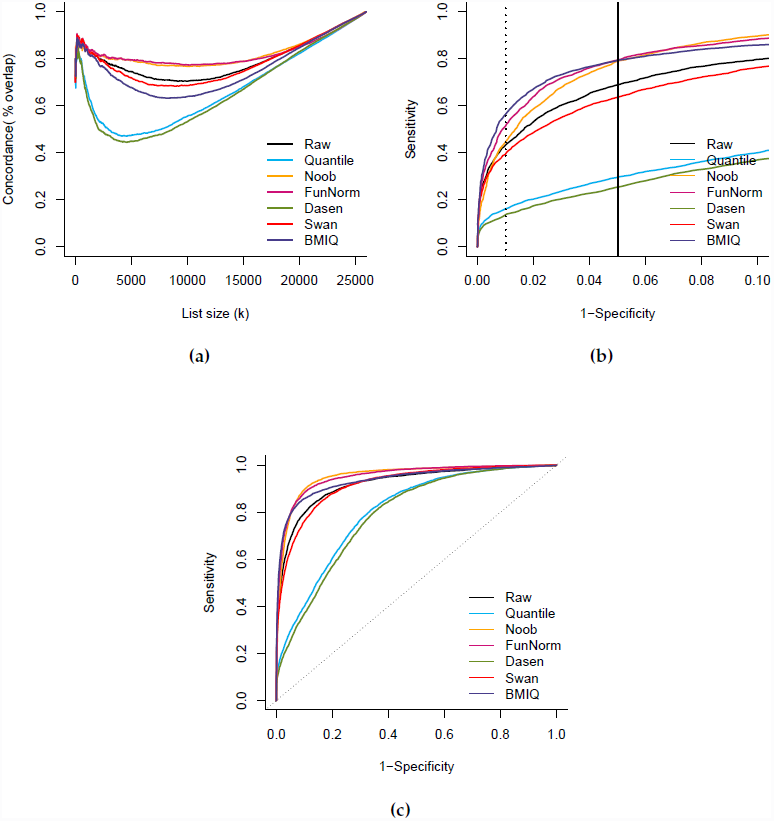
Improvements in replication of tumor subtype heterogeneity. (a) Like Figure 4a (b) Like Figure 4b but for all normalization methods. (c) Like (b) but for the full range of specificity

**Supplementary Figure S7.**
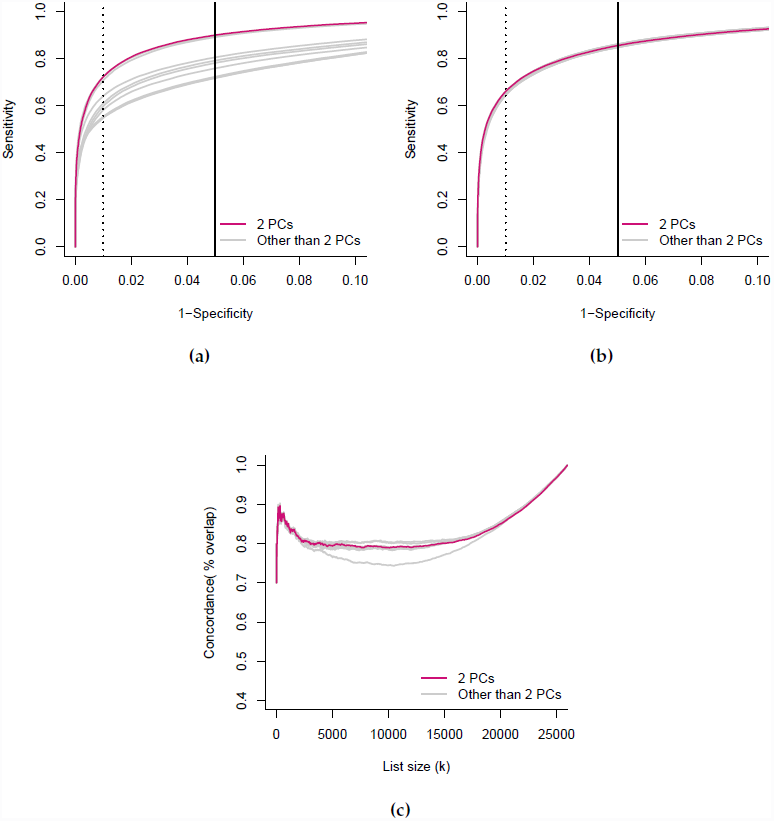
The impact of the number of principal components. (a) Like Figure 2a, (b) Like Figure 3a (c) Like Figure 4a, but showing the difference between using *m* = 2 components and other choices of *m* ≤ 10.

**Supplementary Figure S8.**
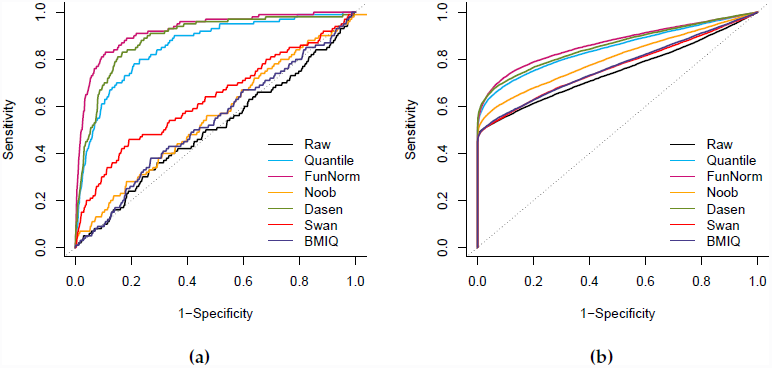
Improvements in blood samples, all methods. Like Figure 5, but for all normalization methods we assess.

## Supplementary material

In the following text we describe exhaustively the transformations that we applied to the control probes to create the control probe summaries that were used throughout the paper as covariates in the functional normalization. Note that we chose the transformations by considering the recommendations made in the GenomeStudio Methylation manual [Ill, 2010].

- For “Bisulfite Conversion I” probes, 3 probes (C1,C2,C3) are expected to have high signal in the green channel in case the bisulfite conversion reaction was successful, and similarly 3 additional probes (C4,C5,C6) are expected to have high signal in the red channel. We therefore consider these 6 intensities and take the mean as a single summary value.
- For “Bisulfite Conversion II” probes, 4 probes are expected to have high intensities in the red channel in case the bisulfite conversion reaction was successful. Therefore we consider the mean of these 4 intensities as a single summary value.
- For the “Extension” control probes, 2 probes must be monitored in the red channel (A,T) and 2 probes must be monitored in the green channel (C,G). We consider the 4 raw intensities as output values for a total of 4 summary values.
- For the “Hybridization” probes, the 3 probes have to be monitored in the green channel. We consider the raw intensities as output values, corresponding to low, medium and high hybridization signals, for a total of 3 summary values.
- For the “Staining” probes, we select the green intensity of the probe that is expected to have high intensity in the green channel, and similarly for the probe that is expected to have high intensity in the red channel. This results in 2 summary values.
- For “Non-polymorphic” controls, we consider the 2 probes that are expected to be high in the green channel (C and G) and the two probes that are expected to be high in the red channel (A and T). We consider the 4 raw intensities as output for a total of 4 summary values.
- “Target removal” probes have to be monitored only in the green channel. We use the raw intensities for the 2 probes as output values for a total of 2 summary values.
- For “Specificity II” probes, we monitor the 3 probes in both the green and red channels, for a total of 6 output values. The green channel is expected to be low and the red channel to be high, so we also form the ration of the mean of the green channel to the mean of the red channel as output value. This results in 7 summary values.
- For “Specificity I” probes, 3 probes are expected to have high signal in the green channel, and 3 different probes are expected to have high signal in the red channel. For each trio, we take the raw intensities in the corresponding “good” channel, giving us 6 output values. For each trio, we also consider the signal-to-noise ratio by taking the ratio of the mean of the 3 intensities measured in the good channel (high signal expected) and the mean of the three intensities in the opposite channel (low signal expected). This gives us 2 ratio summaries. We also consider the mean of these two ratios, giving us 1 additional output value. This results in a total of 9 summary values.
- “Normalization” probes: probes targeting A bases (32) and T bases (61) have to be monitored in the red channel and probes targeting G (32) and C (61) have to be monitored in the green channel. For each type (A,C,T,G), we consider the mean of the intensities in their corresponding channel, that we denote by *normA, normC, normT* and *normG* (4 output values). Moreover, we consider the ratio (*normC + normG*) / (*normA + normT*) as a surrogate for dye bias computed with positive controls (1 additional output value), for a total of 5 summary values.
- For the Out-of-band probes (Oob), we first take the 1st, 50th and 99th percentiles of the 92,596 green intensities (3 output values). Because the variation seen in the green Oob probes is similar to the that of the red Oob probes, we omit the latter. Nevertheless, we consider the ratio of the median of the 92,596 green intensities and the median of the 178,406 red intensities, as a surrogate for dye bias. This results in a total of 4 summary values.

